# Development of Cell Permeable NanoBRET Probes for the Measurement of PLK1 Target Engagement in Live Cells

**DOI:** 10.1101/2023.02.25.529946

**Authors:** Xuan Yang, Jeffery L. Smith, Michael T. Beck, Jennifer M. Wilkinson, Ani Michaud, James D. Vasta, Matthew B. Robers, Timothy M. Willson

## Abstract

PLK1 is a protein kinase that regulates mitosis and is both an important oncology drug target and a potential anti-target of drugs for the DNA damage response pathway or anti-infective host kinases. To expand the range of live cell NanoBRET target engagement assays to include PLK1 we developed an energy transfer probe based on the anilino-tetrahydropteridine chemotype found in several selective PLK inhibitors. Probe **11** was used to configure NanoBRET target engagement assays for PLK1, PLK2, and PLK3 and measure the potency of several known PLK inhibitors. In cell target engagement for PLK1 was in good agreement with the reported cellular potency for inhibition of cell proliferation. Probe **11** enabled investigation of the promiscuity of adavosertib, which had been described as a dual PLK1/WEE1 inhibitor in biochemical assays. Live cell target engagement analysis of adavosertib by NanoBRET demonstrated PLK activity at micromolar concentrations but only selective engagement of WEE1 at clinically relevant doses.

**Graphical Abstract:** 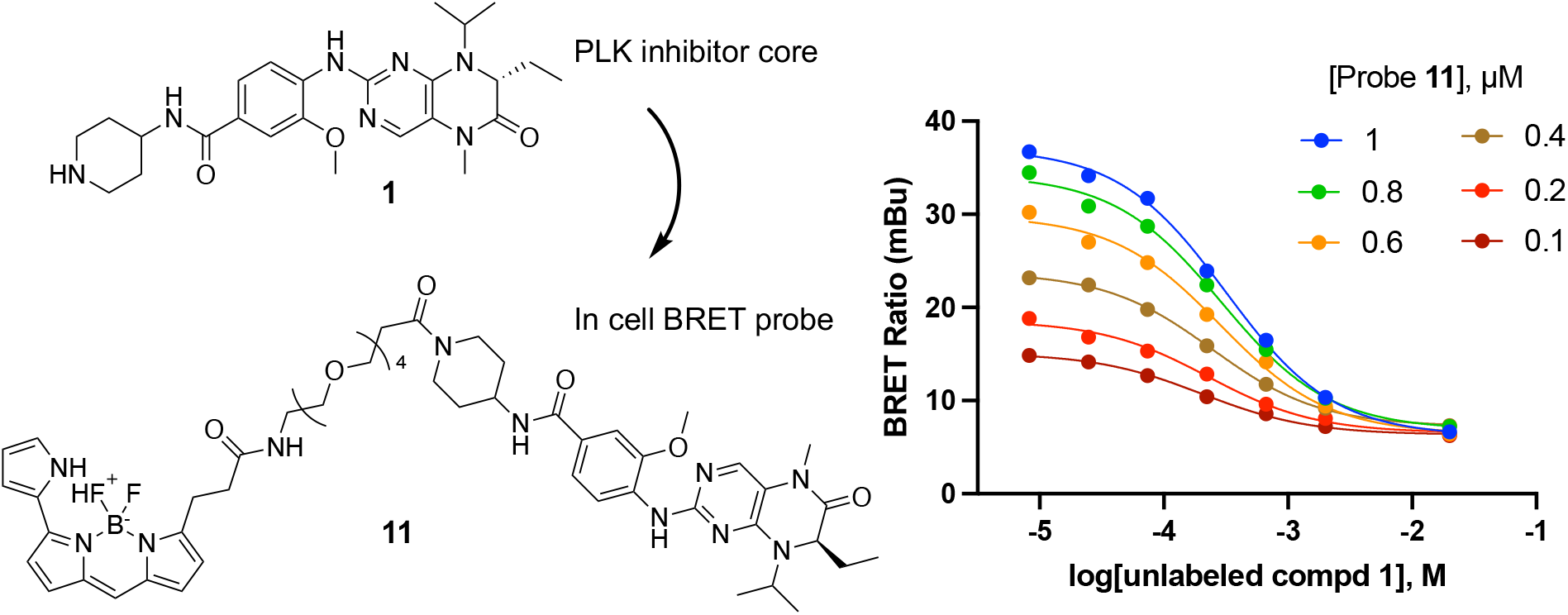

## 1. Introduction

The polo-like kinases are Ser/Thr protein kinases that play key roles in the cell cycle and mitosis and are often dysregulated in cancer [1,2]. Five paralogs are found in humans, PLK1–5, of which PLK1 is the most highly studied member. Inhibition of PLK1 results in aberrant chromosome segregation, mitotic block, and cell death [2]. Notably, PLK2, PLK3, and PLK4 have also been implicated in various aspects of cell cycle control [3,4]. PLK5 lacks a functional kinase domain and is included in the family only because of homology in its other domains. Many ATP-competitive PLK inhibitors have been characterized in biochemical enzyme inhibition assays, with inhibitors of PLK1 often showing activity on PLK2 and PLK3 but little or no activity on PLK4. Conversely, PLK4 inhibitors generally do not show activity on PLK1–3. Although PLK1 inhibitors show robust anticancer activity in animal models, they have been plagued by dose-limiting toxicity in clinical trials and none have been approved as cancer drugs to date [1]. BI2436 was the first PLK1 inhibitor to enter clinical trials. Both BI2436 and its follow on analog volasertib (Figure 1) were reported to be nanomolar inhibitors of PLK1–3 [5,6]. Later generation compounds such as GSK461364 [7] and onvansertib [8] were reported as nanomolar inhibitors with selectivity for PLK1 over PLK2 and PLK3. It remains unclear, however, whether pan-PLK or selective PLK1 inhibition will be clinically superior for treating solid tumors, since there is often a disconnect between the potency of enzyme inhibition and the high micromolar doses required to see therapeutic effects [2].

**Figure 1.**
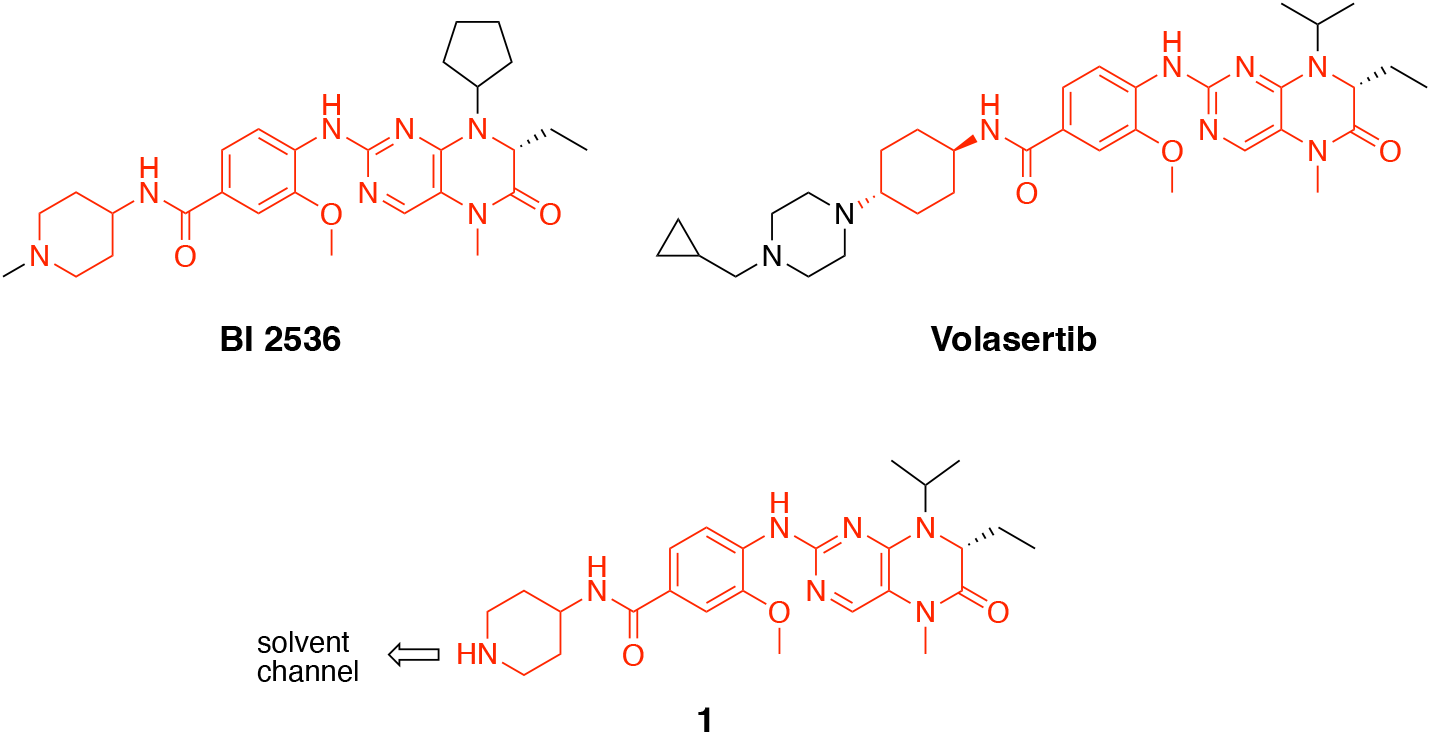
Structures of PLK inhibitors BI 2536 and volasertib, and design of intermediate **1**. The common anilino-tetrahydropteridine core is shown in red.

Beyond their established role as cancer targets, protein kinases have emerged as promising host cell targets for many infectious diseases. For example, viruses often coopt host kinases to effect cell entry and replication [9,10]. Protein kinases also play important roles in the host immune response to viral and bacterial infection, such that kinase inhibitors are being identified for use in combination with conventional anti-infective drugs [11,12]. In addition, suppression of the inflammatory pathways with JAK/STAT inhibitors can aid the recovery of patients in the post-infection phase of the disease [9]. PLK1 has also been reported as a collateral target of inhibitors of WEE1 kinase, a synthetic lethality target of the DNA damage response [13]. In all these cases, PLK1 can be considered as an anti-target since its inhibition results in potential pathway antagonism or can confound optimization of host cell kinase inhibitors for infectious disease therapy due to cell toxicity [14].

We have shown that live cell NanoBRET target engagement assays can be used to measure the cellular potency of protein kinase inhibitors [15,16]. These NanoBRET assays use promiscuous kinase inhibitors conjugated to BODIPY 576/589 as cell penetrant probes (a.k.a. tracers) to generate bioluminescence resonance energy transfer (BRET) when combine with ultrabright NanoLuc (NLuc)-kinase fusion proteins [17]. Competition of the fluorescent tracer by ATP-competitive kinase inhibitors gives a measure of the in-cell target engagement that often correlates with the cellular potency of the inhibitors determined by more time-consuming phospho-substrate assays [15]. Unfortunately, however, none of the promiscuous tracers that have been previously developed were shown to work with PLK1 [15]. Therefore, to address this critical gap in the NanoBRET probe set, we developed a specific tracer for this important signaling kinase starting from a known PLK inhibitor chemotype.

## 2. Results

### 2.1. Synthesis of Fluorescence Energy Transfer Probes 10 and 11

Bifunctional PLK1 energy transfer probes were developed from the anilino-tetrahydropteridine core present in the ATP-competitive PLK inhibitors BI 2536 and volasertib [5,6] (Figure 1). Analysis of the PLK1 crystal structure (PDB: 2RKU) suggested that installation of a linker on the piperidine ring of intermediate **1** would be optimal for placement of the BODIPY 576/589 fluorophore. Synthesis of intermediate **1** started by HATU-mediated coupling between tert-butyl 4-aminopiperidine-1-carboxylate (**2**) and 3-methoxy-4-nitrobenzoic acid (**3**) followed by reduction of the product **4** using hydrazine hydrate solution catalyzed by Raney-Nickel to yield the known aniline **5** [18] (Scheme I). Coupling of aniline **5** with the commercially available tetrahydropteridine **6** using a Buchwald-Hartwig amination gave compound **7**, which was *N*-Boc deprotected under acidic conditions to yield the key intermediate **1**. Two potential energy transfer probes were synthesized from intermediate **1**, first by coupling of intermediate **1** to BODIPY 576/589 NHS ester (**9**) in the presence of DIPEA to yield probe **10**. Alternatively, a tetraethylene glycol (PEG_4_) linker was attached to intermediate **1** followed by coupling to BODIPY 576/589 to yield energy transfer probe **11** (Scheme I).

**Scheme I.**
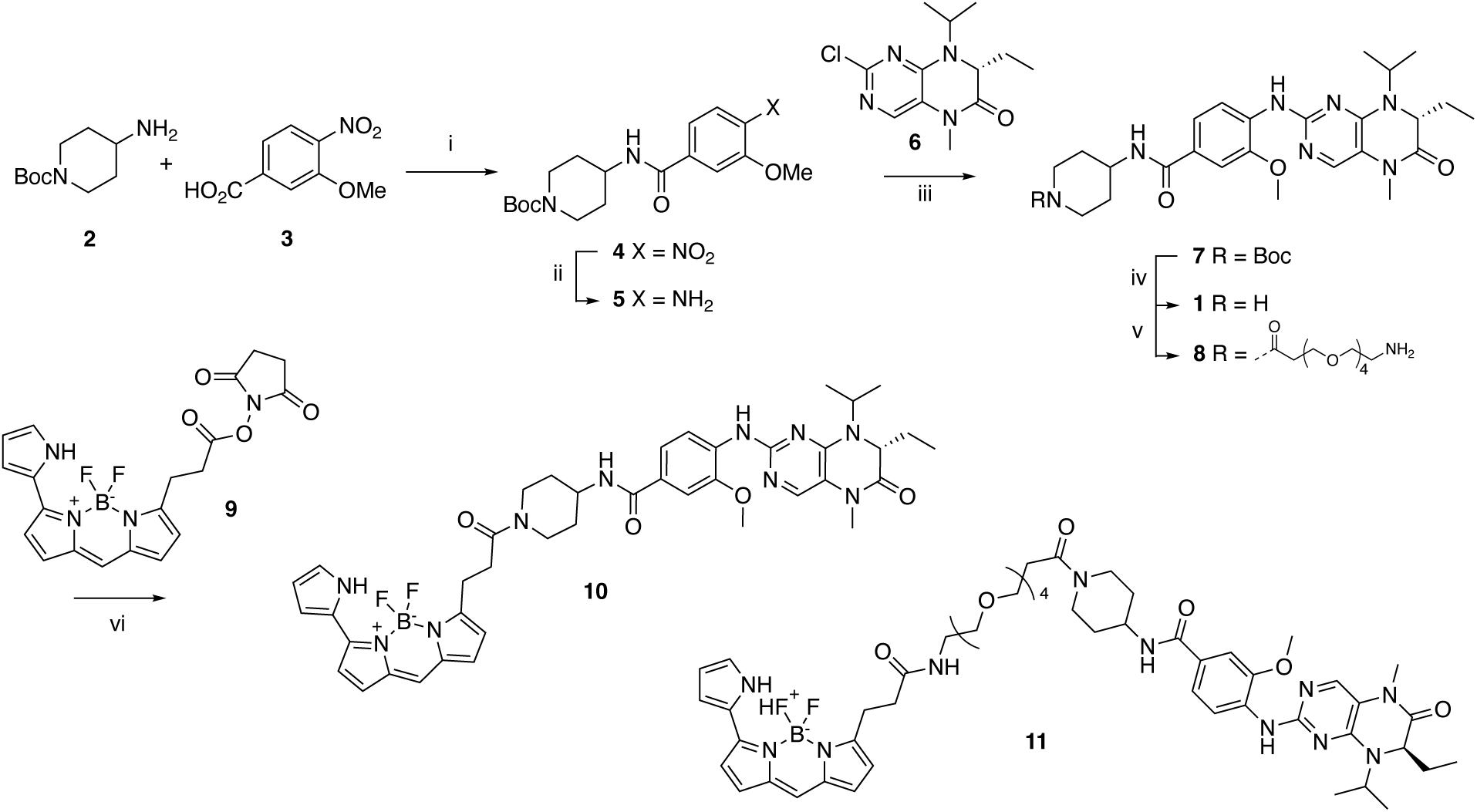
Synthesis route of energy transfer probes **10** and **11**. Reagents and conditions: (i) HATU, DIPEA, DMF, RT, 69%; (ii) hydrazine hydrate solution (50-60%), Raney-Nickel, EtOH, 50 °C, 97%; (iii) K_2_CO_3_, Pd_2_(dba)_3_, XPhos, t-BuOH, 120 °C, microwave irradiation; (iv) TFA, DCM, RT, 59% for two steps; (v) t-Boc-*N*-amido-(PEG)_4_-acetic acid, TBTU, DIPEA, DMF, RT, 58%; then TFA, DCM, RT, 86%; (vi) BODIPY 576/589 NHS ester (**9**) DIPEA, DMF, RT, 68% for energy transfer probe **10**, 70% for energy transfer probes **11**.

### 2.2. Development of a PLK1 NanoBRET assay

We conducted the probe titration studies in live HEK293T cells in the adherent (ADH) format [15] via energy transfer using plasmid DNAs encoding *N*- and *C*-terminal fusions of full-length PLK1 with NLuc. We found that for both probes **10** and **11**, NLuc fused to the *N*-terminus of human PLK1 (NLuc-PLK1) gave a much stronger BRET signal (BRET ratio ∼39 for probe **10** and ∼48 for probe **11**) compared to the NLuc fused to the C-terminus of PLK1 (PLK1-NLuc) (BRET ratios ∼15 for probe **10** and **11**) (Figure 3a). Although both probes gave good BRET ratios using NLuc-PLK1, the signal using probe **11** with the PEG_4_ linker was consistently larger than with probe **10**. Thus, we selected probe **11** for further characterization and optimization of the PLK1 NanoBRET assay. The apparent affinity of intracellular energy transfer probe **11** was measured by treating the cells with increasing concentrations in the presence or absence of an excess (20 μM) of unlabeled intermediate **1** as a competitive PLK1 inhibitor (Figure 2b). The BRET ratio was plotted versus NanoBRET energy transfer probe **11** concentration to determine an apparent intracellular potency, EC_50_ = 0.30 μM. To optimize concentrations for the PLK1 NanoBRET assay the apparent cellular affinity of the unlabeled intermediate **1** was measured at multiple fixed concentrations of the energy transfer probe **11** (Figure 2c) with the goal of selecting a concentration that was at or below the EC_50_ of the probe for the target protein as prior results have demonstrated that lower probe concentrations resulted in more accurate estimation of intracellular compound affinity.

**Figure 2.**
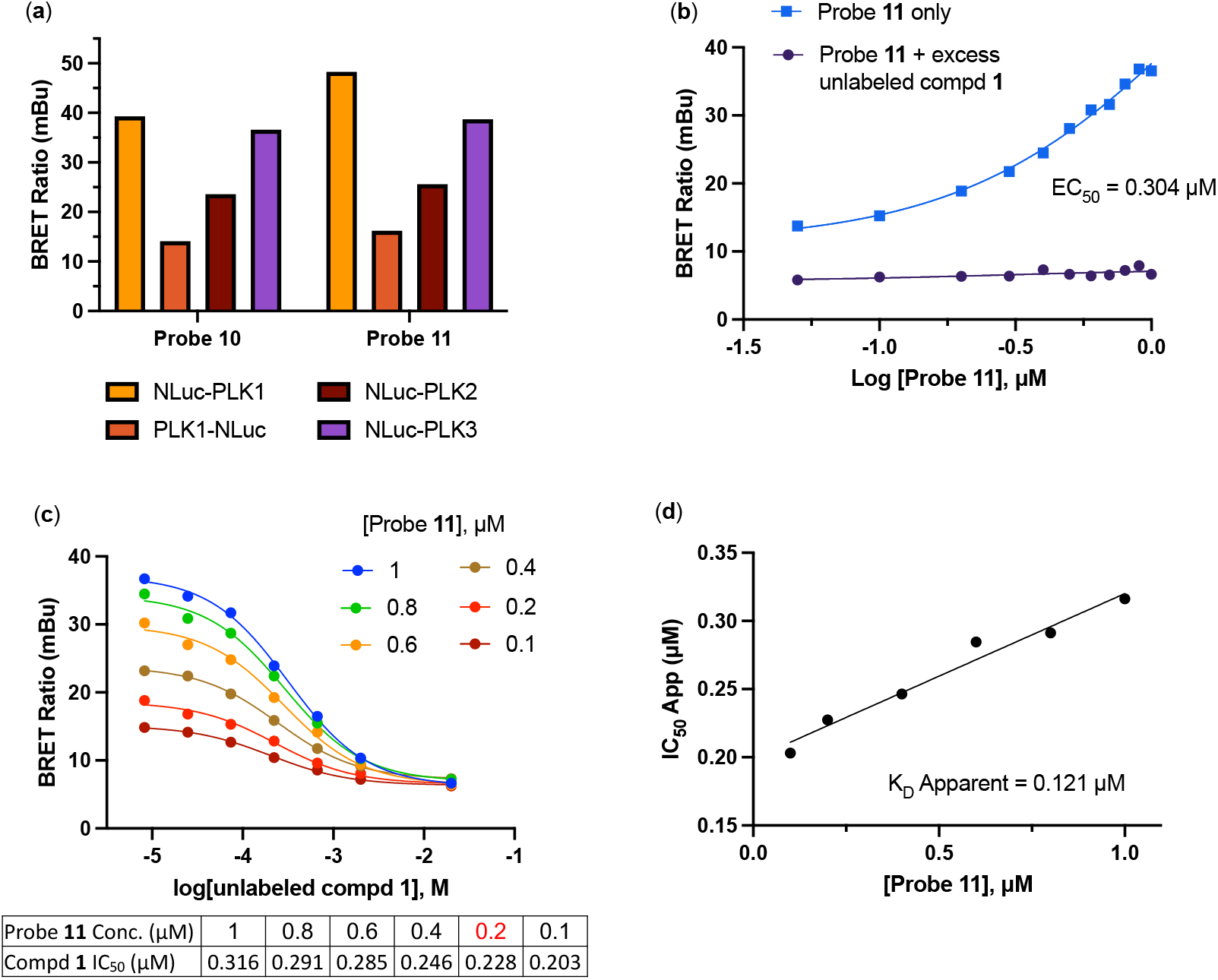
**(a)** BRET ratios observed with 1 µM of probes **10** and **11** on the PLK1, PLK2, and PLK3 NLuc fusion proteins. **(b)** Apparent intracellular NanoBRET energy transfer probe (**11**) affinity for NLuc-PLK1 fusion protein in the adherent (ADH) format. **(c)** Optimization of NanoBRET transfer probe (**11**) concentrations for NLuc-PLK1 fusion protein in the ADH format. **(d)** Quantitative analysis of BRET using Cheng-Prusoff relationship for PLK1 kinase in live HEK293T cells with NanoBRET energy transfer probe **11**. The apparent K_D_ value for compound **1** was determined from the y-intercept by linear regression.

**Figure 3.**
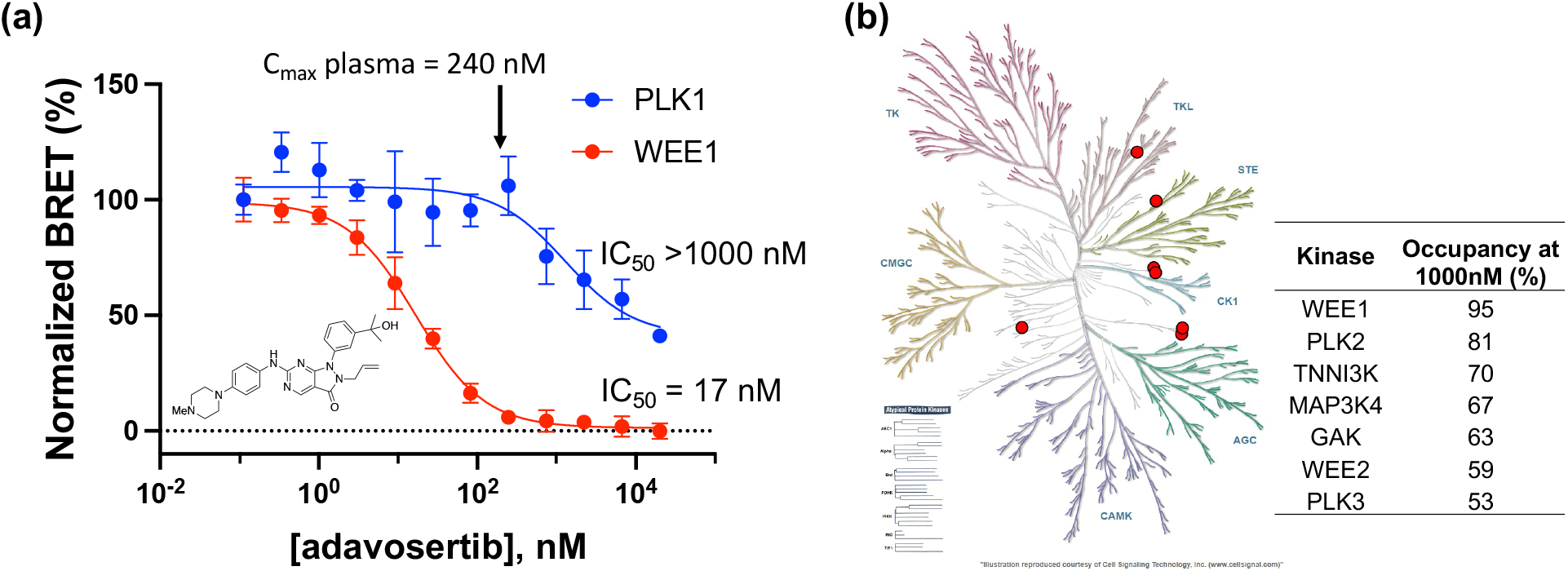
**(a)** Dose response of adavosertib in the PLK1 and WEE1 NanoBRET assays. The IC_50_ for in cell target engagement predicted that only WEE1 would be inhibited at the clinical dose. **(b)** Live cell target occupancy of adavosertib at 1000 nM across 192 Kinase NanoBRET assays using the K10 tracer [24]. Only 7 kinases showed >50% occupancy, shown as the red dots on a kinome tree with the % occupancy noted.

A concentration of 0.2 μM of energy transfer probe **11** was selected as the optimal concentration for the PLK1 NanoBRET assay as it yielded a high assay window (>3 fold) and was below the EC_50_ for the probe. Replotting the apparent IC_50_ values for unlabeled intermediate **1** against the concentration of NanoBRET energy transfer probe **11**, a linearized Cheng-Prusoff analysis yielded an apparent K_D_ of 0.12 μM for unlabeled compound **1** for intracellular target engagement of PLK1 (Figure 2d). To test whether energy transfer probes **10** or **11** could also be used to develop NanoBRET assays for PLK2 and PLK2, the BRET signal was measured using NLuc fused to the *N*-terminal of human PLK2 (NLuc-PLK2) or human PLK2 (NLuc-PLK3). As shown in Figure 1a, robust signals with BRET ratios of 24–38 were obtained using both probes. Since the PLK1 NanoBRET assays had been optimized with probe **11**, we selected the same probe for development of PLK2 and PLK3 NanoBRET assays using the identical procedures. The optimized concentration of energy transfer probe **11** for the NanoBRET assays was 1 μM with both NLuc-PLK2 and NLuc-PLK3 (SI Figure 1), which gave assay windows of 2.4 and 2.1-fold respectively.

### 2.3. In Cell Target Engagement of PLK Inhibitors

Having developed NanoBRET assays for PLK1–3 using energy transfer probe **11**, we selected a series of known PLK inhibitors to compare their enzyme inhibition potency with intracellular target engagement. Several factors can affect the cellular potency of ATP-competitive kinase inhibitors, including their ability to cross cell membranes and the K_m_(ATP) of the individual kinase, which determines the effect of intracellular ATP as a competitor. The IC_50_ for seven PLK inhibitors were determined in the PLK1, PLK2, and PLK3 NanoBRET assays in HEK293T cells. As shown in Table 1, all seven inhibitors demonstrated potent intracellular target engagement on PLK1 and some showed activity on PLK2 and PLK3. The NanoBRET/Enzyme IC_50_ showed that many of the PLK1 inhibitors were less potent in cells although the range differed for each inhibitor (Table 1). Notably, the potency of PLK1 in-cell target engagement was in good agreement with the lowest reported potency for inhibition of cell proliferation.

**Table 1.** Intracellular target engagement potency of known PLK inhibitors

| Compound | Ref. | Enzyme Inhibition<br>IC<sub>50</sub> (nM)<sup>a</sup> |  |  | NanoBRET target<br>engagement IC<sub>50</sub> (nM)<sup>c</sup> |  |  | Cell<br>proliferation<br>IC<sub>50</sub> (nM)<sup>a</sup> |
| --- | --- | --- | --- | --- | --- | --- | --- | --- |
|  |  | PLK1 | PLK2 | PLK3 | PLK1 | PLK2 | PLK3 |  |
| BI2536 | [6] | 0.83 | 3.5 | 9.0 | 3.9 | 9.0 | 42 | 2–25 |
| Volasertib | [5] | 0.87 | 5.0 | 56 | 18 | 23 | 59 | 11–37 |
| TPKI-26 | [14,19] | 2.0 | n.d. | n.d. | 18 | 64 | 1700 | 5 |
| Ro3280 | [20] | 3.0 | n.d. | n.d. | 3.5 | 45 | 180 | 6–45 |
| GSK461364 | [7] | 0.5<sup>b</sup> | 860<sup>b</sup> | 1000<sup>b</sup> | 4.2 | >10000 | >10000 | 10–100 |
| MLN0905 | [21] | 2.0 | n.d. | n.d. | 14 | 6600 | 3000 | 9 |
| Onvansertib | [8] | 2.0 | >10000 | >10000 | 21 | >10000 | >10000 | 42 |

<sup>a</sup>IC<sub>50</sub> values for PLK enzyme inhibition and cell proliferation from the cited references. <sup>b</sup>Time-dependent enzyme inhibition. K<sub>i</sub>(app) from Ref [7]. <sup>c</sup>All NanoBRET data is an average of n = 2.

### 2.4. In Cell Target Selectivity of Adavosertib

To further demonstrate the utility of the PLK1 NanoBRET assay we profiled adavosertib (MK-1775/AZD1775), a WEE1 kinase inhibitor in clinical development for treatment of pancreatic cancer whose kinase selectivity has been the subject of debate in the literature [13,22,23]. In the WEE1 NanoBRET assay using the K10 tracer [24] adavosertib demonstrated an IC_50_ = 17 nM, which is in agreement with the results of inhibition of substrate phosphorylation in cells [23]. In contrast, using the energy transfer probe **11** in the PLK1 NanoBRET assay adavosertib showed >50% target occupancy only at micromolar concentrations, well above its reported circulating plasma levels in patients (Figure 3a). To further annotate the in-cell selectivity of adavosertib we utilized our broad spectrum profiling protocol employing the energy transfer probe K10, which enables assay of 192 kinases but not PLK1 using a single tracer [24]. At a concentration of 1000 nM adavosertib showed >50% target engagement of only seven kinases (Figure 3b). Notably, at this concentration, PLK2 and PLK3 showed 95% and 53% target engagement, respectively.

## 3. Discussion

We have developed an energy transfer probe **11** that can be used to configure NanoBRET assays for PLK1, PLK2, and PLK3, addressing a critical gap in PLK family target engagement profiling. The optimized PLK1 NanoBRET assay allows, for the first time, researchers to determine the intracellular target engagement of PLK1 inhibitors, which further expands the utility of this assay technology to support lead optimization and counter-screening of this important drug target.

Since the cellular concentration of ATP (usually 1–10 mM) is often much higher than the biochemical K_m_(ATP) of kinase enzymes, ATP-competitive kinase inhibitors are often less potent in intact cells than predicted from cell free assay and it is important to determine intracellular target engagement to be able to interpret their phenotypic effects [25]. For PLK1 the biochemical K_m_(ATP) = 2–5 μM [26] and is similar for PLK2 and PLK3. By screening a series of PLK inhibitors in preclinical and clinical development, we demonstrated that their in-cell target engagement was often lower than predicted by their biochemical enzyme inhibition potency (Table 1). While Ro3280 showed no drop off in cellular potency, volasertib was >20-fold less potent in cells than its enzyme potency. The other PLK inhibitors showed a loss in cell potency ranging from 5–11-fold. Each of the seven PLK1 inhibitors have been reported to show cytotoxic activity in cancer cell lines. It is notable that the lowest IC_50_ values for inhibition of cell proliferation closely matched the IC_50_ for intracellular target engagement of PLK1 by NanoBRET for all seven inhibitors (Table 1). BI2536 and volasertib, first and second generation PLK inhibitors in the anilino-tetrahydropteridine chemotype, were also active in the PLK2 and PLK3 NanoBRET assays matching their pan-PLK enzyme inhibition profile. TPKI-26, another derivative of the anilino-tetrahydropteridine chemotype [19], was a dual PLK1/PLK2 inhibitor in the NanoBRET assay but showed only weak activity on PLK3. Ro3280, which was published as a selective PLK1 inhibitor [20], profiled as a pan-PLK inhibitor in cells, but with a preference for PLK1 > PLK2 > PLK3. GSK461324, a time-dependent PLK inhibitor with a complex selectivity profile using FRET-based biochemical assays [7], was shown to be highly PLK1 selective when profiled in the live cell NanoBRET target engagement assay. The third generation inhibitor MLN0905 was also selective for PLK1 in the NanoBRET target engagement assays, although it showed weak activity on both PLK2 and PLK3 which is consistent with its binding to both of these kinases at 1 μM in an Ambit KINOMEscan assay [21]. Finally, the third generation PLK1 inhibitor onvansertib was also highly PLK1 selective in the NanoBRET assay with an IC_50_ on PLK1 that matched its potency for inhibition of cell proliferation [8]. Our results demonstrate that these NanoBRET assays allow, for the first time, direct comparison of the intracellular potency and isozyme selectivity of PLK inhibitors under development as cancer drugs.

The utility of the PLK1 NanoBRET assay to support the profiling of a kinase inhibitor of clinical importance was further applied to the drug adavosertib, which was originally reported as a select WEE1 inhibitor [22] but was subsequently described as a dual WEE1/PLK1 inhibitor on the basis of chemical proteomic and biochemical enzyme inhibition assays [13]. A more recent report suggested that adavosertib was a selective WEE1 inhibitor in cells at clinically relevant doses by profiling the phosphorylation of downstream substrates using immunoblots [23]. These conflicting results opened the possibility that either the biochemical analysis of PLK1 inhibition may not translate in vivo, or that protein phosphorylation analysis may not accurately capture specific PLK1 activity in cells. Using the PLK1 and WEE1 NanoBRET assays combined with broad profiling across 192 kinases, we demonstrated that adavosertib was indeed a selective WEE1 inhibitor in cells at its clinical dose but did also exhibit off target PLK activity at micromolar concentrations with the rank order of PLK2 > PLK3 ∼ PLK1.

In summary, we have developed energy transfer probes to enable a PLK NanoBRET assays that further expand the utility of this technology to profile the in-cell target engagement of kinase inhibitors. These NanoBRET assays provide a robust and rapid method to determine the cellular potency and isozyme selectivity of PLK inhibitors, which can aid lead optimization and counter screening for undesirable off target activity.

## 4. Materials and Methods

### 4.1. Chemistry

#### 4.1.1. General Methods

All chemicals were commercially available except those whose synthesis is described. All reaction mixtures and column eluents were monitored by analytical thin-layer chromatography (TLC) performed on pre-coated Sorbtech fluorescent silica gel plates, 200 μm with an F254 indicator; visualization was accomplished by UV light (254/365 nm). Column chromatography was undertaken with a Biotage Isolera One instrument. Nuclear magnetic resonance (NMR) spectrometry was run on a Varian Inova 400 MHz or Bruker Avance III HD 850 MHz spectrometer. The NMR data were processed using MNova 14.3.1.

Chemical shifts are reported in ppm with residual solvent peaks referenced as internal standard. Analytical LC/MS data were obtained using a Waters Acquity Ultrahigh-performance liquid chromatography (UPLC) system equipped with a photodiode array (PDA) detector running an acetonitrile/water gradient.

#### 4.1.2. Synthesis of Florescence Energy Transfer Probes 10 and 14

Tert-butyl 4-(3-methoxy-4-nitrobenzamido)piperidine-1-carboxylate (**4**)

To a solution of 3-methoxy-4-nitrobenzoic acid (**3**) (1.05 g, 1 Eq, 5.32 mmol) in DMF (12 mL) was added HATU (2.56 g, 1.5 Eq, 7.98 mmol) and DIPEA (1.38 g, 1.9 mL, 2 Eq, 10.64 mmol). The reaction mixture was stirred for 10 min at room temperature before adding tert-butyl 4-aminopiperidine-1-carboxylate (**2**) (1.06 g, 1 Eq, 5.32 mmol). After 12 hours, the reaction mixture was concentrated, and the resulting material was purified by chromatography on silica gel (eluent of ethyl acetate / hexane = 0-40%) to give the impure product. The desired product was recrystallized in the test tubes (40% ethyl acetate in hexanes). The white crystal was filtered and dried under vacuum to afford the desired product (**4**) (1.40 g, 69 %). TLC condition: Hexanes/ethyl acetate = 1/1. ^1^H NMR (400 MHz, CD_3_OD) δ 7.86 (d, *J* = 8.2 Hz, 1H), 7.69 (s, 1H), 7.50 (d, *J* = 8.9 Hz, 1H), 4.18 – 4.04 (m, 3H), 4.03 (s, 3H), 3.02 – 2.85 (m, 2H), 2.01 – 1.93 (m, 2H), 1.59 – 1.49 (m, 2H), 1.49 – 1.46 (m, 9H). ^13^C NMR (101 MHz, CD_3_OD) δ 167.5, 156.5, 153.6, 143.0, 140.9, 126.1, 120.3, 114.0, 81.2, 57.3, 49.1, 32.5, 28.7.

Tert-butyl 4-(4-amino-3-methoxybenzamido)piperidine-1-carboxylate (**5**)

To a suspension of tert-butyl 4-(3-methoxy-4-nitrobenzamido)piperidine-1-carboxylate (**4**) (900 mg, 1 Eq, 2.37 mmol) in Ethanol (20 mL) was added hydrazine hydrate solution (50-60%) (1.7 mL) and the reaction mixture was stirred at 50 °C for 15 min until the starting material is dissolved completely. Excess of Raney-Nickel (1.39 g, 10 Eq, 23.7 mmol) was added to the reaction mixture, and the reaction was maintained at 50 °C. After 1 h when gas evolution ceased, Raney-Nickel was filtered under vacuum. The filtrate was concentrated in vacuo to give the desired product (**5**) (806 mg, 97 %). TLC condition: hexanes/ethyl acetate = 2/1. ^1^H NMR (400 MHz, CD_3_OD) δ 7.36 – 7.26 (m, 2H), 6.73 – 6.67 (m, 1H), 4.11 (d, *J* = 13.8 Hz, 2H), 4.06 – 3.99 (m, 1H), 3.89 (s, 3H), 3.01 – 2.79 (m, 2H), 1.91 (dd, *J* = 12.6, 3.4 Hz, 2H), 1.57 – 1.43 (m, 11H). ^13^C NMR (101 MHz, CD_3_OD) δ 169.8, 156.5, 147.8, 142.5, 123.8, 122.3, 114.2, 110.5, 81.1, 56.1, 48.6, 32.8, 28.7. MS ESI *m/z*: 350.1 [M+H]^+^.

(*R*)-4-((7-ethyl-8-isopropyl-5-methyl-6-oxo-5,6,7,8-tetrahydropteridin-2-yl)amino)-3-methoxy-N-(piperidin-4-yl)benzamide (**1**)

Tert-butyl 4-(4-amino-3-methoxybenzamido)piperidine-1-carboxylate (**5**) (600 mg, 1 Eq, 1.72 mmol), (*R*)-2-chloro-7-ethyl-8-isopropyl-7,8-dihydropteridin-6(5*H*)-one (**6**) (437 mg, 1 Eq, 1.72 mmol), 2-(dicyclohexylphosphanyl)-2’,4’,6’-tris(isopropyl)biphenyl (XPhos) (164 mg, 0.2 Eq, 343 µmol), K_2_CO_3_ (949 mg, 4 Eq, 6.87 mmol) and Pd_2_(dba)_3_ (157 mg, 0.1 Eq, 172 µmol) were added in t-BuOH (15 mL) under Ar atmosphere. The reaction mixture was degassed and purged with Ar. It was then heated in a microwave at 120 °C for 30 min. The mixture was filtered through Celite, washed with DCM and concentrated. The resulting material was purified by chromatography on silica gel (methanol/DCM = 0 - 10%) to give the crude product (**7**), 0.91 g, as a light-yellow solid. This crude product (**7**) was used for the next step directly. TLC condition: DCM/MeOH = 10/1. MS ESI *m/z*: 582.2 [M+H]^+^.

To a solution of crude compound **7** (300 mg) in DCM (3 mL), TFA (3 mL) was added at room temperature. The reaction mixture was stirred for 1 hour at room temperature. The reaction solution was concentrated to give a crude product which was purified with C18 column (0-20% ACN in water + 0.1% TFA) to afford the desired product (**1**) as trifluoroacetate salt (193 mg, 332 μmol, 59% for two steps). It was converted to HCl salt by adding 1M HCl in diethyl ether and concentrated. TLC condition: DCM/MeOH = 10/1+ 5% NH_4_OH. ^1^H NMR (850 MHz, CD_3_OD) δ 8.02 (d, *J* = 7.6 Hz, 1H), 7.66 (s, 1H), 7.65 (s, 1H), 7.61 (d, *J* = 7.5 Hz, 1H), 4.63 – 4.54 (m, 2H), 4.24 – 4.19 (m, 1H), 4.03 (s, 3H), 3.52 (d, *J* = 12.5 Hz, 2H), 3.32 (s, 3H), 3.21 – 3.15 (m, 2H), 2.24 – 2.20 (m, 2H), 2.14 – 2.08 (m, 1H), 2.01 – 1.94 (m, 3H), 1.48 (dd, *J* = 16.1, 6.2 Hz, 6H), 0.88 (t, *J* = 7.2 Hz, 3H). ^13^C NMR (214 MHz, CD_3_OD) δ 168.8, 164.5, 154.4, 152.4, 150.4, 133.0, 129.8, 124.1, 123.8, 121.2, 117.8, 111.6, 61.2, 56.9, 53.3, 49.5, 49.4, 46.5, 44.5, 29.5, 28.9, 28.9, 20.6, 19.5, 8.4. MS ESI *m/z*: 482.3 [M+H]^+^.

(*R*)-*N*-(1-(3-(5,5-difluoro-7-(1H-pyrrol-2-yl)-5H-5^λ^,6^λ^-dipyrrolo[1,2-c:2’,1’-*f*][1,3,2]diazaborinin-3- yl)propanoyl)piperidin-4-yl)-4-((7-ethyl-8-isopropyl-5-methyl-6-oxo-5,6,7,8-tetrahydropteridin-2- yl)amino)-3-methoxybenzamide (**10**)

(*R*)-4-(4-((7-ethyl-8-isopropyl-5-methyl-6-oxo-5,6,7,8-tetrahydropteridin-2-yl)amino)-3- methoxybenzamido)piperidin-1-ium, trifluoracetate (**1**) (11 mg, 1 Eq, 18 μmol) was charged into a flask and was taken up in anhydrous DMF (0.5 mL). To the stirred solution was added DIPEA (12 mg, 16 μL, 5 Eq, 92 μmol), then BODIPY™ 576/589 NHS ester (**9**) (7.9 mg, 1 Eq, 18 μmol). The mixture was stirred for 1 hour and then subjected to preparative HPLC (column: Phenomenex, Luna 5 μM Phenyl-Hexyl, 100 Å, 75×30 mm, 5 micron; mobile phase A: water (0.05% TFA), B: methanol; method: 20%- 100% B, 6 min + 100% B, 4 min). The desired product (**10**) was obtained as a dark purple solid (10 mg, 13 μmol, 68 %). TLC condition: DCM/MeOH=10/1. ^1^H NMR (400 MHz, CD_3_OD) δ 7.98 (d, *J* = 8.4 Hz, 1H), 7.59 (s, 1H), 7.58 (d, *J* = 1.9 Hz, 1H), 7.53 (dd, *J* = 8.4, 1.9 Hz, 1H), 7.23 (s, 1H), 7.22 – 7.15 (m, 3H), 7.00 (d, *J* = 4.6 Hz, 1H), 6.92 (d, *J* = 3.9 Hz, 1H), 6.37 – 6.30 (m, 2H), 4.64 – 4.47 (m, 3H), 4.21 – 4.05 (m, 2H), 3.98 (s, 3H), 3.29 – 3.14 (m, 6H), 2.88 – 2.78 (m, 3H), 2.12 – 1.83 (m, 4H), 1.63 – 1.47 (m, 2H), 1.42 (dd, *J* = 9.2, 6.8 Hz, 6H), 0.84 (t, *J* = 7.4 Hz, 3H). ^13^C NMR (214 MHz, CD_3_OD) δ 172.7, 168.6, 164.5, 156.2, 154.4, 152.3, 152.2, 150.5, 139.0, 135.0, 133.3, 133.2, 129.8, 127.4, 126.8, 124.9, 124.7, 124.2, 123.5, 121.1, 121.0, 119.0, 117.7, 117.6, 112.4, 111.5, 61.1, 56.7, 53.2, 49.9 (2C), 46.0, 43.8, 42.3, 33.8, 28.8, 28.7, 25.8, 20.5, 19.4, 8.3. MS ESI *m/z*: 793.3. [M+H]^+^.

(*R*)-*N*-(1-(1-amino-3,6,9,12-tetraoxapentadecan-15-oyl)piperidin-4-yl)-4-((7-ethyl-8-isopropyl-5-methyl-6-oxo-5,6,7,8-tetrahydropteridin-2-yl)amino)-3-methoxybenzamide (**8**)

To a solution of 2,2-dimethyl-4-oxo-3,8,11,14,17-pentaoxa-5-azaicosan-20-oic acid (t-Boc-N-amido-PEG4-acid) (38 mg, 1 Eq, 0.10 mmol) in DMF (2 mL), TBTU (50 mg, 1.5 Eq, 0.16 mmol) and *N*-ethyl-*N*-isopropylpropan-2-amine (54 mg, 4 Eq, 0.42 mmol) were added and stirred for 15 min. (*R*)-4-(4- ((7-ethyl-8-isopropyl-5-methyl-6-oxo-5,6,7,8-tetrahydropteridin-2-yl)amino)-3- methoxybenzamido)piperidin-1-ium, trifluoracetate (**1**) (62 mg, 1 Eq, 0.10 mmol) was added to the reaction mixture and stirred for 18 hours. Water was added to the reaction mixture and extracted with ethyl acetate. The crude product was purified by column chromatography on silica gel (from 100% DCM to DCM/MeOH=10/1) to afford tert-butyl (*R*)-(15-(4-(4-((7-ethyl-8-isopropyl-5-methyl-6-oxo-5,6,7,8- tetrahydropteridin-2-yl)amino)-3-methoxybenzamido)piperidin-1-yl)-15-oxo-3,6,9,12- etraoxapentadecyl)carbamate (50 mg, 60 μmol, 58 %). TLC condition: DCM/MeOH=10/1. ^1^H NMR (850 MHz, CD_3_OD) δ 8.52 (d, 1H), 7.75 (s, 1H), 7.50 – 7.47 (m, 2H), 4.69 (hept, *J* = 6.9 Hz, 1H), 4.60 – 4.55 (m, 1H), 4.32 (dd, *J* = 7.7, 3.5 Hz, 1H), 4.17 – 4.11 (m, 1H), 4.11 – 4.06 (m, 1H), 4.00 (s, 3H), 3.80 – 3.71 (m, 2H), 3.64 – 3.57 (m, 12H), 3.48 (t, *J* = 5.6 Hz, 2H), 3.31 (s, 3H), 3.26 – 3.18 (m, 3H), 2.81 – 2.77 (m, 1H), 2.76 – 2.70 (m, 1H), 2.68 – 2.62 (m, 1H), 2.07 – 2.03 (m, 1H), 2.00 – 1.95 (m, 1H), 1.94 – 1.87 (m, 1H), 1.80 – 1.72 (m, 1H), 1.64 – 1.56 (m, 1H), 1.56 – 1.50 (m, 1H), 1.47 – 1.39 (m, 15H), 0.84 (t, *J* = 7.5 Hz, 3H). ^13^C NMR (101 MHz, CD_3_OD) δ 171.9, 169.1, 165.2, 156.3, 153.3, 148.5, 139.8, 134.8, 127.5, 121.4, 117.4, 117.0, 110.1, 80.0, 71.6, 71.5 (3C), 71.4 (3C), 71.2, 71.0, 68.6, 67.6, 59.4, 56.6, 49.7, 48.7, 46.3, 42.2, 41.3, 34.5, 33.4, 32.4, 28.8, 28.6, 21.5, 20.0, 9.1. MS ESI *m/z*: 829.4 [M+H]^+^

Tert-butyl (*R*)-(15-(4-(4-((7-ethyl-8-isopropyl-5-methyl-6-oxo-5,6,7,8-tetrahydropteridin-2- yl)amino)-3-methoxybenzamido)piperidin-1-yl)-15-oxo-3,6,9,12-tetraoxapentadecyl)carbamate (24 mg, 1 Eq, 29 μmol) was dissolved to the DCM (1.5 mL) before adding TFA (0.56 mL). The reaction mixture was stirred at room temperature for 3 hours and then concentrated to give the desired product (**8**) (21 mg, 25 μmol, 86 %). ^1^H NMR (400 MHz, CD_3_OD) δ 7.90 (d, J = 8.3 Hz, 1H), 7.56 (s, 1H), 7.51 (d, J = 1.9 Hz, 1H), 7.46 (dd, J = 8.3, 1.9 Hz, 1H), 4.54 – 4.41 (m, 3H), 4.14 – 3.94 (m, 2H), 3.90 (s, 3H), 3.74 – 3.64 (m, 4H), 3.63 – 3.52 (m, 12H), 3.22 (s, 3H), 3.21 – 3.11 (m, 1H), 3.06 (t, J = 5.1 Hz, 2H), 2.79 – 2.57 (m, 3H), 2.09 – 1.78 (m, 4H), 1.60 – 1.42 (m, 2H), 1.36 (t, J = 7.0 Hz, 6H), 0.78 (t, J = 7.4 Hz, 3H). MS ESI *m/z*: 729.3. [M+H]^+^.

(*R*)-*N*-(1-(1-(5,5-difluoro-7-(1H-pyrrol-2-yl)-5H-5^λ^,6^λ^-dipyrrolo[1,2-c:2’,1’-*f*][1,3,2]diazaborinin-3-yl)-3- oxo-7,10,13,16-tetraoxa-4-azanonadecan-19-oyl)piperidin-4-yl)-4-((7-ethyl-8-isopropyl-5-methyl-6-oxo- 5,6,7,8-tetrahydropteridin-2-yl)amino)-3-methoxybenzamide (**11**)

(*R*)-15-(4-(4-((7-ethyl-8-isopropyl-5-methyl-6-oxo-5,6,7,8-tetrahydropteridin-2-yl)amino)-3- methoxybenzamido)piperidin-1-yl)-15-oxo-3,6,9,12-tetraoxapentadecan-1-aminium, trifluoracetate (**8**) (12 mg, 1 Eq, 13 μmol) was dissolved in anhydrous DMF (0.4 mL). To the stirred solution was added DIPEA (8.1 mg, 11 μL, 5 Eq, 63 μmol) then BODIPY™ 576/589 NHS ester (**9**) (5.4 mg, 1 Eq, 13 μmol). The reaction mixture was stirred at room temperature for 1 hour and then purified with preparative HPLC (column: Phenomenex, Luna 5 μM Phenyl-Hexyl, 100 Å, 75×30 mm, 5 micron; mobile phase A: water (0.05% TFA), B: methanol; method: 20%-100% B, 6 min + 100% B, 4 min). The desired product (**11**) was obtained as a dark purple solid (9 mg, 9 μmol, 70 %). ^1^H NMR (850 MHz, CD_3_OD) δ 7.99 (dd, *J* = 8.4, 2.3 Hz, 1H), 7.60 – 7.56 (m, 2H), 7.52 (ddd, *J* = 8.4, 4.1, 1.9 Hz, 1H), 7.22 – 7.19 (m, 2H), 7.19 – 7.16 (m, 2H), 7.00 (dd, *J* = 4.5, 2.9 Hz, 1H), 6.90 (dd, *J* = 3.9, 1.8 Hz, 1H), 6.36 – 6.33 (m, 1H), 6.31 (dd, *J* = 4.0, 1.9 Hz, 1H), 4.59 – 4.52 (m, 2H), 4.52 – 4.48 (m, 1H), 4.14 (tt, *J* = 11.3, 4.2 Hz, 1H), 4.08 – 4.04 (m, 1H), 4.00 (s, 3H), 3.77 – 3.69 (m, 2H), 3.64 – 3.57 (m, 12H), 3.54 (t, *J* = 5.5 Hz, 2H), 3.39 – 3.37 (m, 2H), 3.36 (s, 3H), 3.28 – 3.25 (m, 4H), 3.23 – 3.17 (m, 1H), 2.81 – 2.76 (m, 1H), 2.74 – 2.69 (m, 1H), 2.65 – 2.60 (m, 2H), 2.11 – 2.05 (m, 1H), 2.05 – 2.01 (m, 1H), 2.00 – 1.95 (m, 1H), 1.94 – 1.87 (m, 1H), 1.62 – 1.55 (m, 1H), 1.55 – 1.48 (m, 1H), 1.45 – 1.39 (m, 6H), 0.84 (t, *J* = 7.5 Hz, 3H). ^13^C NMR (214 MHz, CD_3_OD) δ 174.8, 172.0, 168.6, 164.4, 156.3, 154.3, 152.1, 150.3, 138.9, 134.9, 133.2, 133.0, 129.8, 127.4, 127.0, 124.9, 124.6, 124.1, 123.2 (2C), 121.0 (2C), 119.1, 117.6, 117.2, 112.3, 111.4, 71.6, 71.5, 71.4, 71.3, 70.6, 68.6, 61.0, 56.8, 53.1, 49.9, 49.5, 49.4, 46.3, 42.2, 40.5, 36.1, 34.5, 33.3, 32.3, 28.9, 28.7, 25.7, 20.6, 19.3, 8.3. MS ESI *m/z*: 1040.4. [M+H]^+^.

### 4.2. Biology

#### 4.2.1. Cell Culture

HEK293 cells (ATCC) were cultured in DMEM (Gibco) + 10% FBS (Seradigm), with incubation in a humidified, 37 °C/5% CO_2_ incubator. N-terminal NLuc-PLK1, PLK2 or PLK3 fusions were encoded in pFN31K expression vectors (Promega). Carrier DNA was encoded in pFN5K vectors (Promega). For cellular BRET target engagement experiments, HEK-293 were transfected with NLuc/target fusion constructs using FuGENE HD (Promega) according to the manufacturer’s protocol. Briefly, NLuc/target fusion constructs were diluted into Transfection Carrier DNA (Promega) at a mass ratio of 1:9 (mass/mass), after which FuGENE HD was added at a ratio of 1:3 (µg DNA: µL FuGENE HD). 1 part (vol) of FuGENE HD complexes thus formed were combined with 20 parts (vol) of HEK-293 cells suspended at a density of 2 × 10^5^ per mL, followed by incubation in a humidified, 37 °C/5% CO_2_ incubator for 20 hr.

#### 4.2.2 NanoBRET assays

In Cell BRET Assays

All BRET assays were performed in white, tissue-culture treated 96-well plates (Corning #3917) using adherent HEK-293 cells at a density of 2 × 10^4^ cells per well. All chemical inhibitors were prepared as concentrated stock solutions in DMSO (Sigma-Aldrich) and diluted in Opti-MEM to prepare working stocks. Cells were equilibrated for 2 h with the appropriate energy transfer probe and test compound prior to BRET measurements. Energy transfer probes were prepared at a working concentration of 20× in Tracer dilution buffer (12.5 mM HEPES, 31.25% PEG-400, pH 7.5). Individual kinase NanoBRET assays used the following energy transfer probes and concentrations: PLK1, probe **11** (0.2 µM); PLK2, probe **11** (1.0 µM); PLK3, probe **11** (1.0 µM); WEE1, tracer K10 (Promega, 0.13 µM). To measure BRET, NanoBRET NanoGlo Substrate and Extracellular NLuc Inhibitor (Promega) were added according to the manufacturer’s recommended protocol, and filtered luminescence was measured on a GloMax Discover luminometer equipped with 450 nm BP filter (donor) and 600 nm LP filter (acceptor), using 0.5 s integration time. BRET ratios are calculated by dividing the acceptor luminescence by the donor luminescence. Milli-BRET (mBRET) units (mBU) are calculated by multiplying the raw BRET ratios by 1,000. Broad spectrum profiling on 192 kinases was performed using the K192 NanoBRET Target Engagement assay (Promega) with tracer K10 using the published protocol [24].

Initial evaluation of BRET probes **10** and **11**

To test BRET probes **10** and **11** against PLK1, HEK293 adherent cells were incubated with either NLuc fused to the N-terminus of PLK1, (NLuc-PLK1) or the C-terminus of PLK1, (PLK1-NLuc) using 1 µM of the BRET probe and milliBRET units were measured. The combination with the highest milliBRET units was selected for further processing. NLuc-PLK2 and NLuc-PLK3 were tested with both probes **10** and **11** using the same protocol.

##### Apparent intracellular potency of BRET probe **11** for NanoLuc-PLK1

To estimate the half-maximal effective concentration (EC_50_) of probe **11** to PLK1 in cells, adherent HEK293 cells in a 96-well plate expressing NLuc-PLK1 fusion protein were mixed with increasing concentration of BRET probe **11**. The resulting BRET ratio was fitted to the sigmoidal, 4-parameter dose-response curve using GraphPad. As a control, the same experiment was repeated in the presence of an excess of unlabeled compound **1** (20 μM) as a competitive inhibitor for 2 hours before adding 3×Complete Substrate plus Inhibitor Solution.

Apparent intracellular potency of compound **1** to PLK1, PLK2 and PLK3 using BRET assay Adherent HEK293 cells in a 96-well plate expressing NLuc-PLK1, NLuc-PLK2 or NLuc-PLK3 fusion proteins were mixed with increasing concentration of the BRET probe **11** and various concentrations of unlabeled compound **1** for 2 hours before adding 3×Complete Substrate plus Inhibitor Solution. For each probe concentration used, the BRET ratios obtained were used to estimate compound **1** half-maximal inhibitory concentrations (IC_50_) by fitting the data to the four-parameter, sigmoidal dose-response curve in GraphPad. To obtain the apparent in-cell, binding potency (K_D_ apparent) of **1** towards the PLK1, we used linear regression of the estimated IC_50_ versus the concentration of the BRET probe.

## Supplementary Materials

Figure S1: Tracer titration of probe **11** on NLuc-PLK2 and NLuc-PLK3. ^1^H and ^13^C NMR Spectra of intermediate **1**, probe **10**, and probe **11**.

## Author Contributions

Conceptualization, T.W., X.Y. and M.R.; Methodology X.Y.; Investigation; X.Y., J.S., M.B., J.V., A.M. and J.W.; Writing – Original Draft Preparation, T.W. and X.Y.; Writing – Review and Editing, M.B. and J.S.

## Acknowledgements

We thank Rafael Couñago (SGC-UNC) for editing the biology methods.

## Sample Availability

The NanoBRET probe **11** and vectors for NLuc-PLK1 and are available from the authors.

