## Supplementary Material for "Development of Cell Permeable NanoBRET Probes for the Measurement of PLK1 Target Engagement in Live Cells"

**Figure S1:** Tracer titration of probe **11** on NLuc-PLK2 and NLuc-PLK3

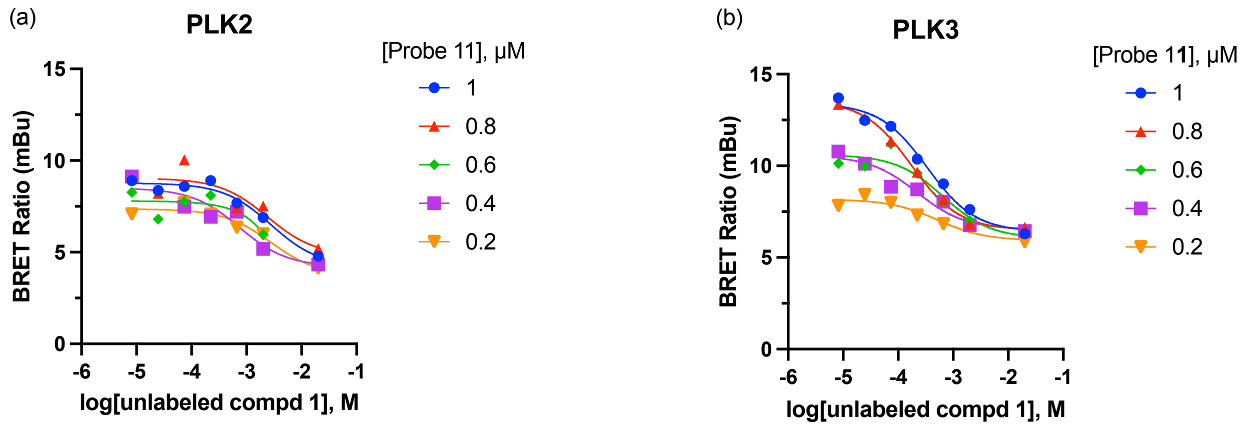


BRET ratios of (a) NLuc-PLK2 and (b) NLuc-PLK3 using multiple concentration of probe **11** with increasing doses of intermediate **1** as a competitor. The 1 μM concentration of probe **11** was selected for use in the PLK2 and PLK3 NanoBRET assays.

$^{1}H$ and $^{13}C$ NMR Spectra of intermediate **1**, probe **10**, and probe **11**:

$^{1}H$ NMR spectrum of intermediate **1** in MeOD

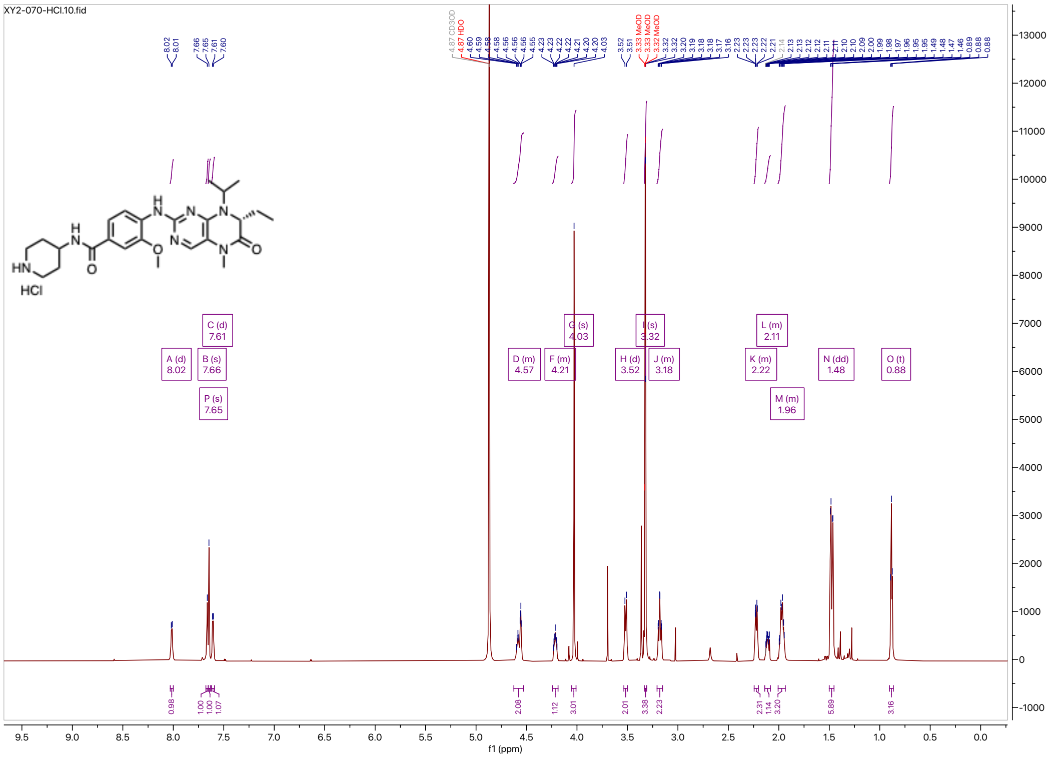


$^{13}C$ NMR spectrum of intermediate **1** in MeOD

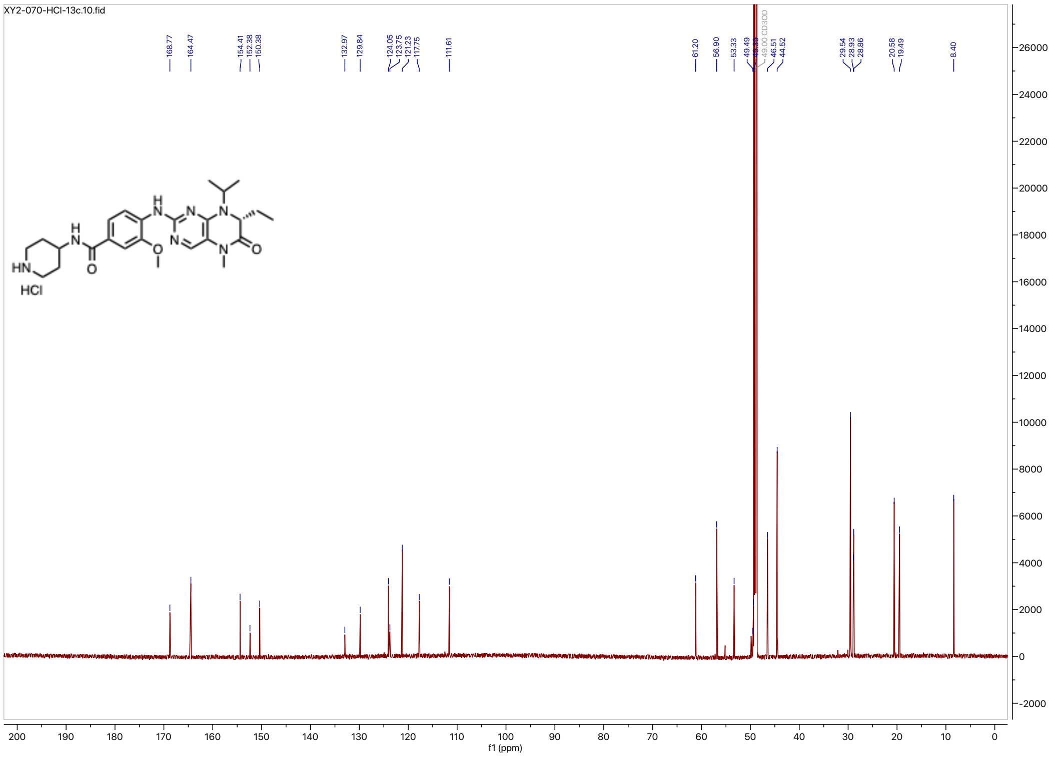

[1]H NMR spectrum of probe **10** in MeOD

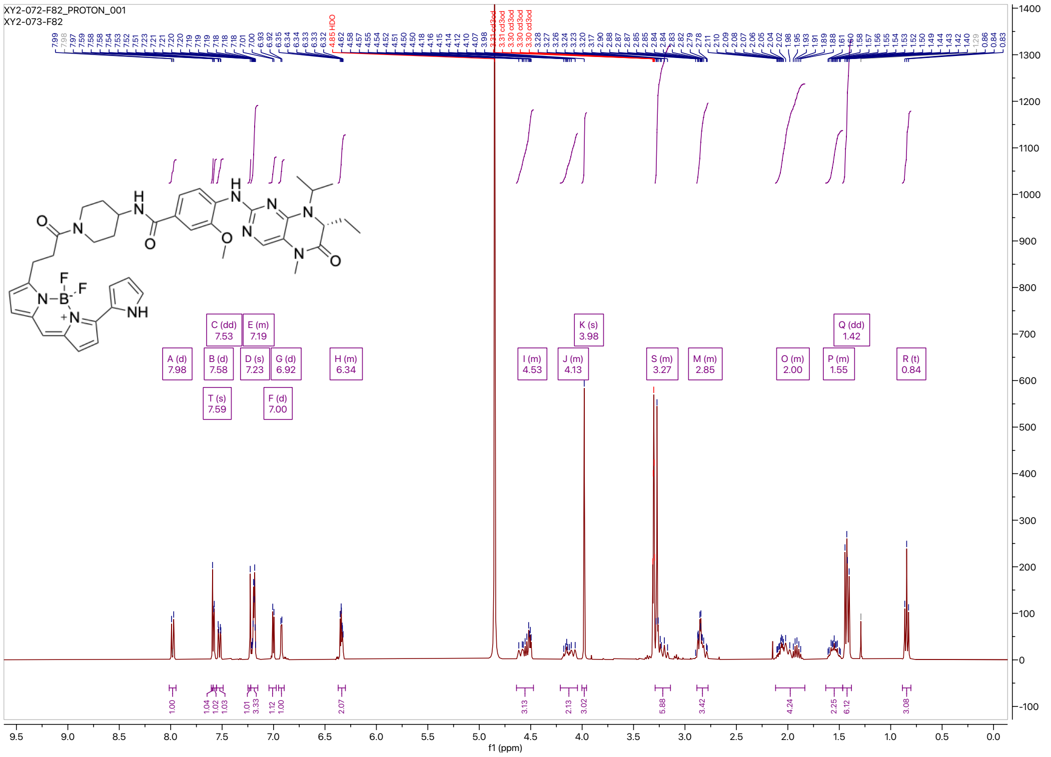

[13]C NMR spectrum of probe **10** in MeOD

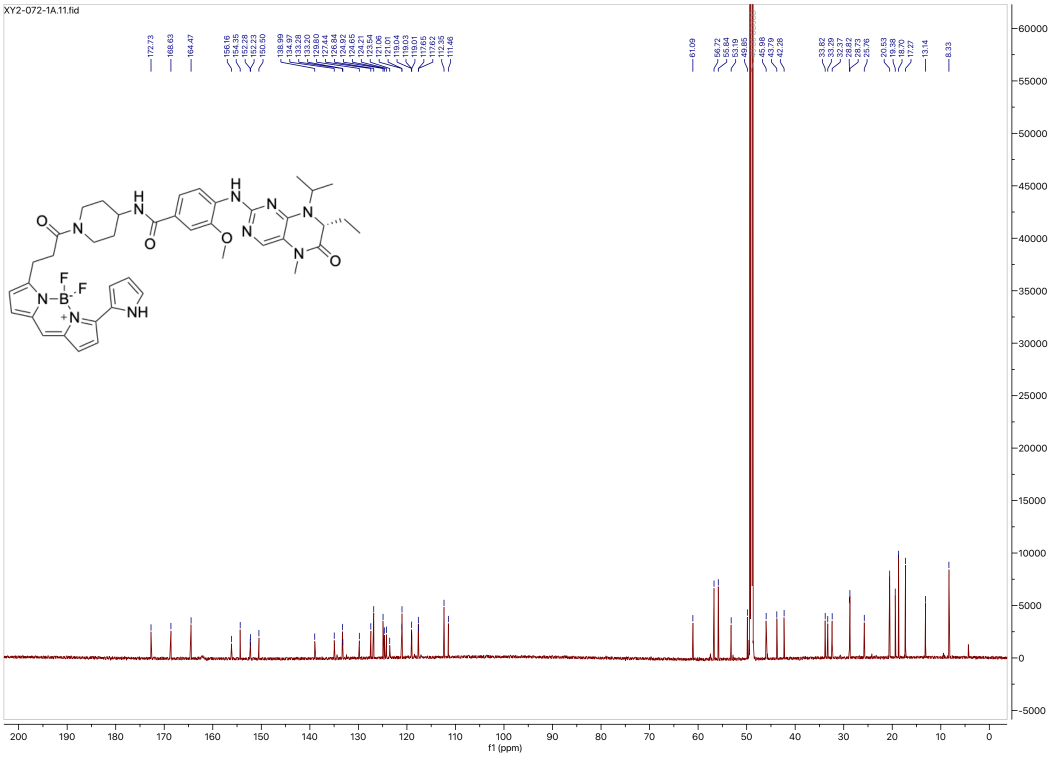

### [1]H NMR spectrum of probe **11** in MeOD

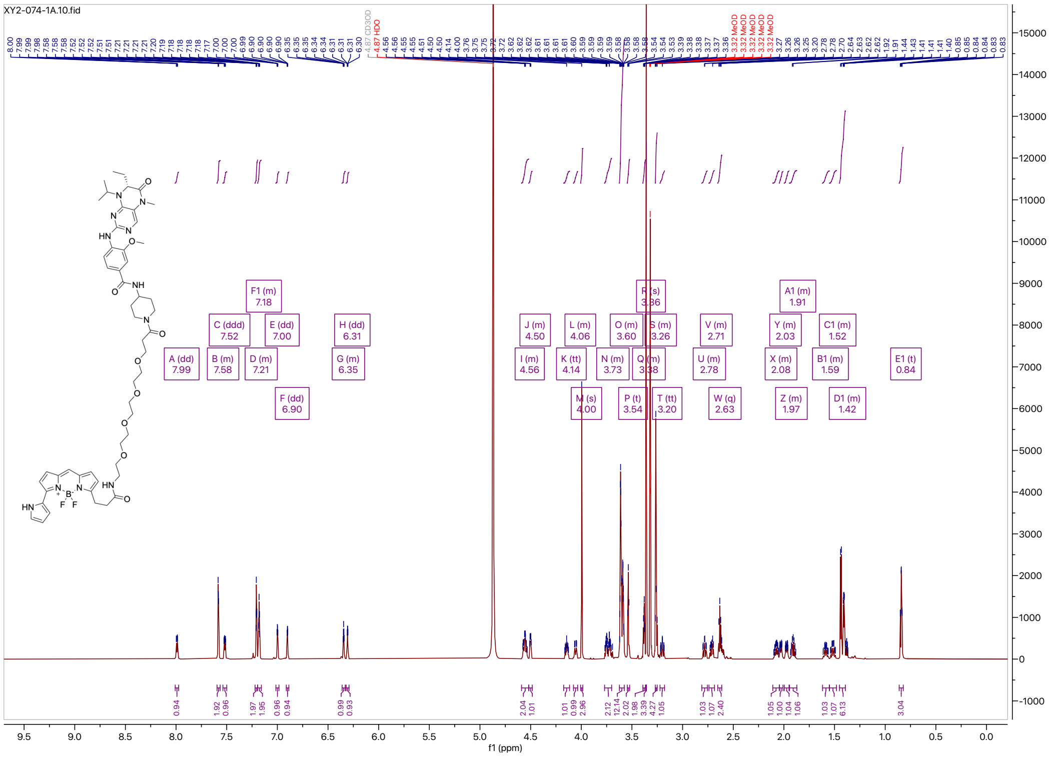


### [13]C NMR spectrum of probe **11** in MeOD

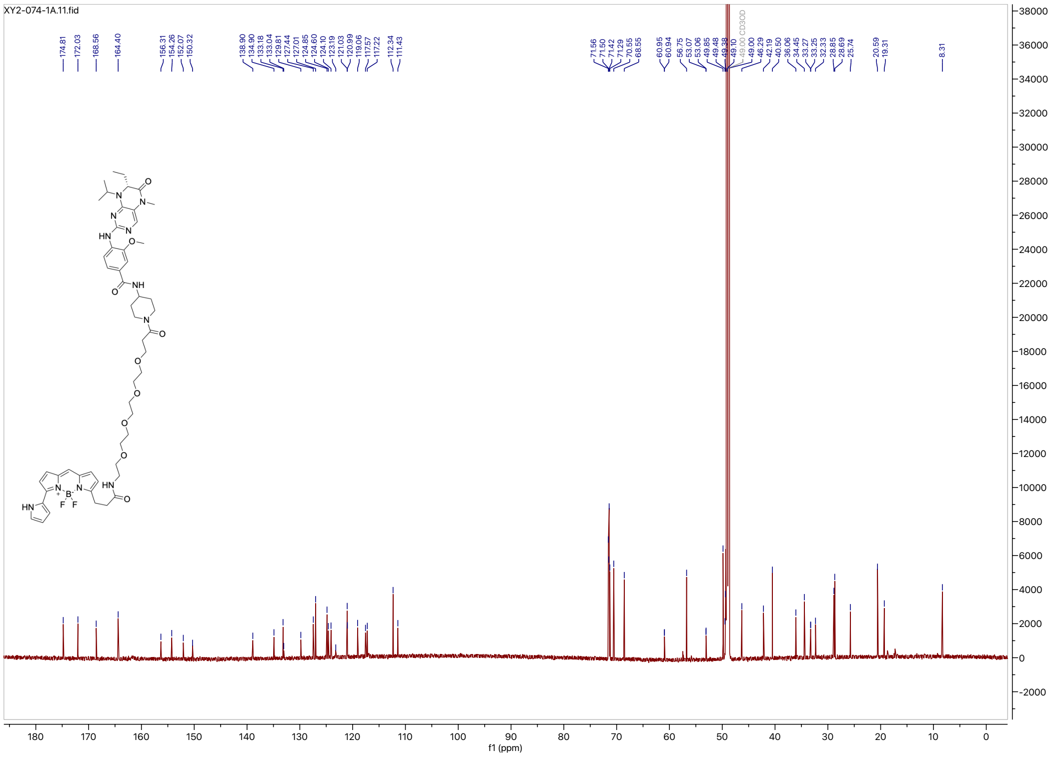

HSQC spectrum of probe **11** in MeOD

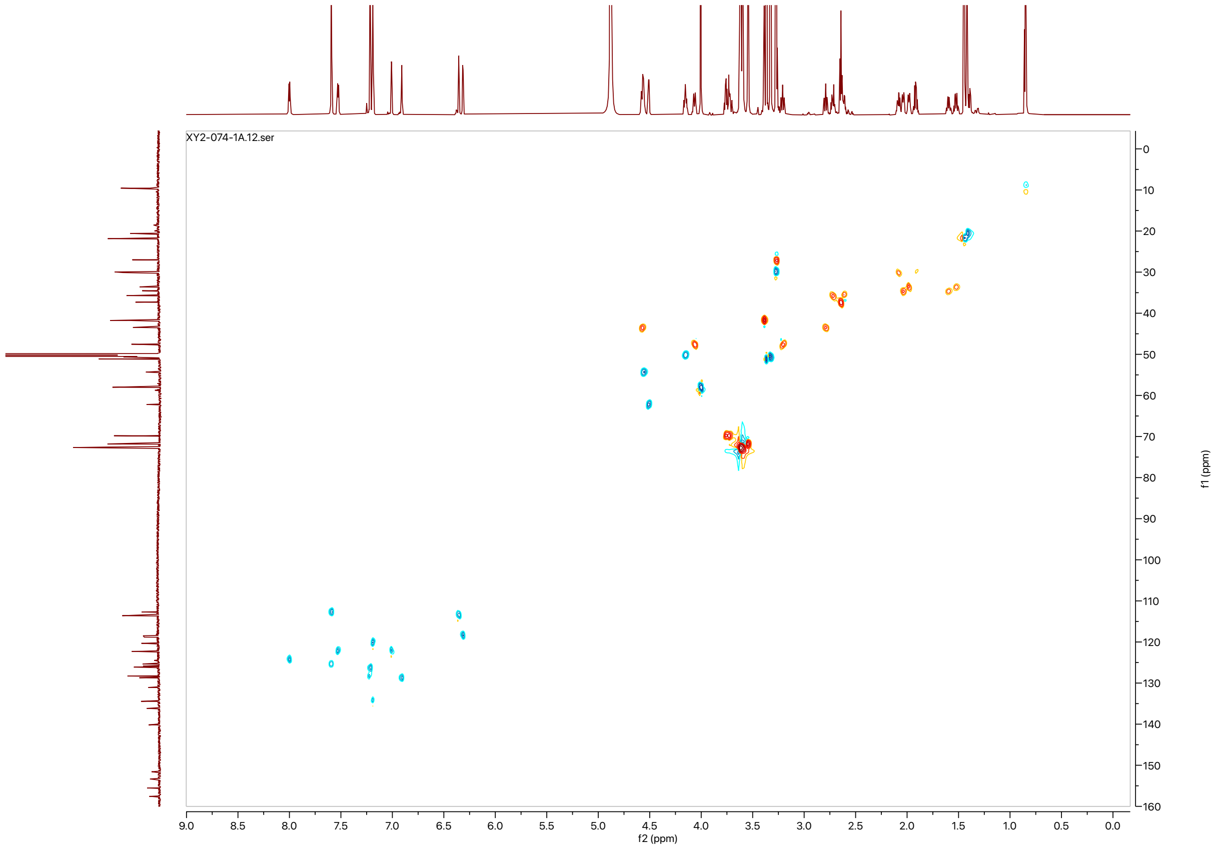
